# Comprehensive evaluation of differential serodiagnosis between Zika and dengue viral infection

**DOI:** 10.1101/421628

**Authors:** Day-Yu Chao, Matthew T. Whitney, Brent S. Davis, Freddy A Medina, Jorge L Munoz, Gwong-Jen J. Chang

**Author notes:** **Corresponding author: Chang, Gwong-Jen J**, or **Day-Yu Chao**.

## Abstract

Diagnostic testing for Zika virus (ZIKV) or dengue virus (DENV) infection can be accomplished by a nucleic acid detection method; however, a negative result does not exclude infection due to the low virus titer during infection depending on the timing of sample collection. Therefore, a ZIKV- or DENV-specific serological assay is essential for the accurate diagnosis of patients and to prevent potential severe health outcomes. A retrospective study design with dual approaches of collecting human serum samples for testing was developed. All serum samples were extensively evaluated by using both non-infectious virus-like particles (VLPs) and soluble non-structural protein 1 (NS1) in the standard immunoglobulin M (IgM) antibody-capture enzyme-linked immunosorbent assay (MAC-ELISA). Both VLP- and NS1-MAC-ELISAs were found to have similar sensitivity for detecting anti-premembrane/envelope and NS1 antibodies from ZIKV-infected patient sera. Group cross reactive (GR)-antibody-ablated homologous fusion peptide-mutated (FP)-VLPs consistently showed higher P/N values than homologous wild-type VLPs. Therefore, FP-VLPs were used to develop the algorithm for differentiating ZIKV from DENV infection. Overall, the sensitivity and specificity of the FP-VLP-MAC-ELISA and the NS1-MAC-ELISA were each higher than 80% with no statistical significance. A novel approach to differentiate ZIKV from DENV infection serologically has been developed. The accuracy can reach up to 95% when combining both VLP and NS1 assays. In comparison to current guidelines using neutralization tests to measure ZIKV antibody, this approach can facilitate laboratory screening for ZIKV infection, especially in regions where DENV infection is endemic and capacity for neutralization testing does not exist.

## Introduction

Zika virus (ZIKV) and dengue virus (DENV), members of the *Flaviviridae* family, are associated with the resurgence of mosquito-transmitted diseases worldwide.^1^ While DENV continues to impose a great economic and public health burden in tropical and subtropical countries, the recent emergence of ZIKV, circulated in Central and South America since 2013, has resulted in terrifying outbreaks with severe health outcomes, including Guillain-Barre syndrome in adults as well as microcephaly, congenital neurologic malformations, and fetal demise in fetuses.^2, 3^ Clinically, ZIKV and DENV share similar symptoms of infection, geographical distribution, and transmission cycles between humans and *Aedes aegypti* mosquitoes.^4^ A confirmatory diagnosis can be attained by virus isolation or viral RNA detection in serum and other body fluids, but given the low virus titer during ZIKV infection and the high incidence of mild or asymptomatic ZIKV infections, a ZIKV-specific serological assay is essential to accurately diagnosis the patients who were negative by virus isolation or viral RNA detection.^5, 6^

Mosquito-borne flaviviruses can be serologically classified into several complexes, including medically important members of the Japanese encephalitis virus (JEV) complex, (DENV), yellow fever virus (YFV), as well as the recently emerged ZIKV.^7^ During natural infection, the majority of elicited antibodies (Abs) recognize structural proteins pre-membrane (prM) and envelope (E), and non-structural protein 1 (NS1).^7, 8, 9^ Anti-E antibodies that recognize all members of the flavivirus group, members from different serocomplexes, or members within a serocomplex, are classified as group-reactive (GR), complex-reactive (CR), or type-specific (TS)-Abs, respectively.^10, 11, 12^ Although GR or CR anti-NS1 antibodies could be found from other flavivirus infections, recent studies suggested the majority of anti-NS1 antibodies from primary ZIKV infections are dominated by TS Abs and can be used as serological markers to differentiate ZIKV from DENV infections.^8, 13^ However, the cross-reactivity of human anti-NS1 antibodies increased after sequential DENV and ZIKV infections.^8^ Furthermore, the low sensitivity in detecting anti-NS1 antibodies and the discrepancy in determining sero-positivity between detecting anti-E and anti-NS1 antibodies were continuously reported.^14, 15^ Serological cross-reactivity between flaviviruses is common and several recent publications have shown the global efforts trying to resolve this issue to determine the status of ZIKV infection.^13, 16, 17, 18^ Although a validated, virus specific sero-diagnostic test is urgently needed, a rigorous evaluation of the assay is required to ensure optimal patient care and accurate epidemiologic surveillance in regions with active transmission of both DENV and ZIKV.

The objectives of this study were to develop (Phase I) and validate (Phase II) a sero-diagnostic assay that can reliably distinguish and diagnose current/acute ZIKV and/or DENV infection in humans. In the Phase I, we selected and applied several well-characterized, archived serum panels, collected during the 2008 West Nile virus outbreak in South Dakota, the 2009 DENV outbreak in Brazil and the 2016 introduction of ZIKV to Puerto Rico, to thoroughly evaluate anti-prM/E and anti-NS1 IgM antibodies against ZIKV and DENV virus-like particles (VLP) and soluble NS1 antigens. We applied the Receiver Operation Characteristic (ROC) analysis to estimate the proper cut-off and to determine an algorithm that can specifically distinguish and diagnose ZIKV and DENV infection using acute/convalescent human serum specimens. We then conducted a double-blind study using clinical serum specimens collected and provided by Division of Vector-borne Disease (DVBD)-Dengue Branch, Centers for Disease Control and Prevention (CDC), in Puerto Rico to validate the reliability of the algorithm developed in Phase I. Using the classical immunoglobulin M (IgM) antibody-capture enzyme-linked immunosorbent assay (MAC-ELISA), we were able to differentiate between ZIKV and DENV with accuracy higher than 85%. Furthermore, combining both VLP and NS1-MAC-ELISAs, 95% accuracy could be achieved.

## Materials and Methods

### Study design and human serum panels

A two-stage retrospective study design was implemented here including a developmental serum panel in Phase I and a validation panel in Phase II. Serum specimens of all suspected DENV- or ZIKV-infected patients were evaluated by real-time reverse-transcription polymerase chain reaction (rRT-PCR) for acute-stage specimens and MAC-ELISA for IgM seroconversion of the convalescent-phase specimens by the CDC Dengue Branch in San Juan, Puerto Rico.^19^ Since there was a possibility of IgM antibody cross-reactivity between closely related flaviviruses from prior flaviviral infections or vaccination, a supplementary focus-reduction micro-neutralization test (FRμNT) was conducted for all specimens positive by MAC-ELISA. FRμNT is still the only reference standard test by far for differentiating flavivirus infection serologically.

The retrospective, archived serum panels (summarized in Table 1) were used as developmental panels in Phase I to establish the proper cut-off value for the assay and algorithm. Only the ZIKV-infected patient serum panel (the testing panel, Table S3) was collected in Puerto Rico after the first confirmation of ZIKV circulation in the Americas; the rest of the archived specimens were collected prior to the first appearance of ZIKV in the Americas. The ZIKV patient serum panel used for testing included forty-two acute and convalescent ZIKV-infected patient serum pairs from Puerto Rico in 2016, confirmed by rRT-PCR and CDC MAC-ELISA. Acute specimens were collected within seven days and the convalescent phase specimens were taken within seven to thirty days after onset of symptoms. The ninety percent end-point FRμNT (FRμNT_90_) on ZIKV and DENV-2 was used to verify recent infections by following the CDC guidelines.^20^ Primary ZIKV infection is determined by an anti-DENV-2 FRμNT_90_ titer of less than 20 from both acute and convalescent sera with a concurrent positive ZIKV titer (≥20). Secondary ZIKV infection is defined by a positive ZIKV titer with an anti-DENV-2 titer of equal to or greater than 20 from either acute or convalescent sera.

**Table 1.** Characteristics of serum panels used in this study.

| Serum Panel | No of<br>specimens | Single/paired<sup>#</sup> | RT-PCR | 90%FR<sub>μ</sub>NT | Country | Year |
| --- | --- | --- | --- | --- | --- | --- |
| **Testing panel** |  |  |  |  |  |  |
| ZIKV | 42 | Paired | (+) | 4-fold increase | Puerto Rico | 2016 |
| DENV | 54 | Paired | (+) | 4-fold increase | Brazil | 2009 |
| WNV | 97 | Single | n/d<sup>^</sup> | Highest titer | South Dakota, USA | 2008 |
| Other* | 76 | Single |  |  |  |  |

A double-blinded test of VLP- and NS1-MAC-ELISA were conducted in Phase II to validate the established diagnostic algorithm (Table S5). Serum specimens used in this Phase were all collected from Puerto Rico based on convenient series, with only single serum collection from each participant. This study was conducted and reported in accordance with the Standards for Reporting Diagnostic Accuracy Studies (STARD) guidelines.^21^ Informed consent documents for all eligible participants were waived based on the protocol number 6874. An institutional review board (IRB) waiver to use this serum panel for research purposes was approved by the CDC-human studies review board. Since all the specimens were de-identified, the basic demographic and clinical characteristics of the participants were not available in this study.

### Plasmid construction, soluble protein expression and antibody production

A transcriptional and translational optimized eukaryotic cell expression plasmid was used as the backbone to express NS1 protein or pre-membrane/envelope (prM/E) that generate VLPs from ZIKV BPH-2016 strain (Brazil 2016) based on standard molecular cloning procedures as described previously.^22, 23^ The constructed plasmids were electroporated into COS-1 cells using a protocol described previously. VLPs and soluble nonstructural protein 1 (sNS1) were expressed by COS-1 cells electroporated with recombinant expression plasmids encoding the prM/E and NS1 genes, respectively. Electroporated cells were recovered in 150 cm^2^ culture flasks with 50 ml DMEM and incubated at 28°C with 5% CO_2_ for VLP/sNS1 expression. VLPs and sNS1 from DENV-2 strain 16681, DENV-3 strain C0331/94, and West Nile virus (WNV) strain NY99 were produced as described previously^22, 23^ and used in this study. Additionally, the prM/E expressing plasmid was modified by site-directed mutagenesis of an epitope recognized by GR antibody^24^ and the VLPs generated were named ZIKV-FP-VLP or DENV2-FP-VLP.

Both the anti-ZIKV polyclonal rabbit and mouse sera containing high-titer immunoglobulin recognizing all potential antigenic epitopes were generated at the Centers for Disease Control and Prevention, Fort Collins, U.S.A. (US-CDC). Anti-DENV-2, DENV-3, or WNV VLP, and anti-NS1 polyclonal rabbit sera were produced in-house as described previously. Murine hyperimmune ascetic fluid (MHIAF) specific for DENV-2 or DENV-3, or WNV were obtained from the Diagnostic and Reference Laboratory, DVBD-CDC.

### VLP- and NS1-specific MAC/GAC-ELISAs

Human serum specimens were assayed for the presence of prM/E- and NS1-specific antibodies using MAC-ELISAs as previously described.^14, 20, 22, 23^ Briefly, 96-well plates were coated with goat anti-human IgM or IgG (Kirkegaard & Perry Laboratories, Gaithersburg, MD) diluted 1:2,000 in PBS at pH 9.0 and incubated at 4°C overnight. The infected patient serum as well as negative control serum were diluted 1:1,000 in wash buffer (PBS with 0.05% Tween-20), 50 μl were added to wells, and incubated at 37°C for 60 min. ZIKV, DENV, WNV VLPs and NS1, pre-determined and standardized at an optical density of 1.0 at 450nm (OD_450_).by antigen-capture ELISA (Ag-ELISA), were diluted in wash buffer and tested against each serum sample in triplicate.

To deplete anti-prM/E antibodies from serum samples, Ag-ELISA was used to capture VLP immune-complexes in 96-well plates as previously suggested.^22, 23^ In brief, the patient and negative control sera were diluted 1:1,000 in PBS, mixed with VLP antigens, and added immediately to wells pre-coated with anti-prM/E rabbit sera and incubated at 37°C for 60 min. A total of 50 μL of prM/E antibody-depleted sera were transferred to 96-well plates pre-coated with anti-human IgM for performing the NS1-specific MAC-ELISA as described above.

### Data processing and statistical analysis

Both positive and negative values were determined as the average OD_450_ from triplicate samples of each specimen (P) or normal human control sera (N) reacting with VLP or NS1 antigens, respectively. Positive-to-negative (P/N) ratios were derived for each specimen as well as positive and negative serum controls on each plate for validation of the quality of the assay. The P/N ratios from the ZIKV patient sera were compared with the ratios from different serum specimens from the test panel and the positive likelihood ratio (LR+), shown as ROC curve, was calculated by dividing sensitivity by 1-specificity to determine the optimal cutoff value of P/N ratios from VLP- and NS1-MAC-ELISAs.

The Bland-Altman plot was used to measure the consistency of higher P/N values of the ZIKV-over DENV2-FP-MAC-ELISA, or ZIKV-over DENV2-NS1-MAC-ELISA by plotting the ratios of the two methods’ P/N ratio values (ratio of P/N value between ZIKV-MAC-ELISA and DENV2-MAC-ELISA) versus the averages of P/N values from both methods. Two-by-two contingency tables were prepared to determine the sensitivity, specificity, positive predictive value (PPV) and negative predictive value (NPV) of the assays based on the algorithm generated in this study according to the validation serum panel. For all statistical analyses, we used GraphPad Prism version 6 and *p* values less than 0.05 were considered statistically significant.

## Results

### Participant serum panels

Figure 1 showed the flowchart of the archived serum panel retrospectively collected for Phase I and Phase II study. Since the first index patient reported onset of symptoms on November 23, 2015, Puerto Rico became the first U.S jurisdiction to report local transmission of ZIKV when DENV was already endemic in Puerto Rico. In order to proper establishment of the cut-off value of the assay, the criteria of all ZIKV and DENV-confirmed specimens used in Phase I included paired sera collected during the acute and convalescent phase of illness and confirmation of disease status by FRμNT_90_ on ZIKV and DENV-2.

**Fig 1.**
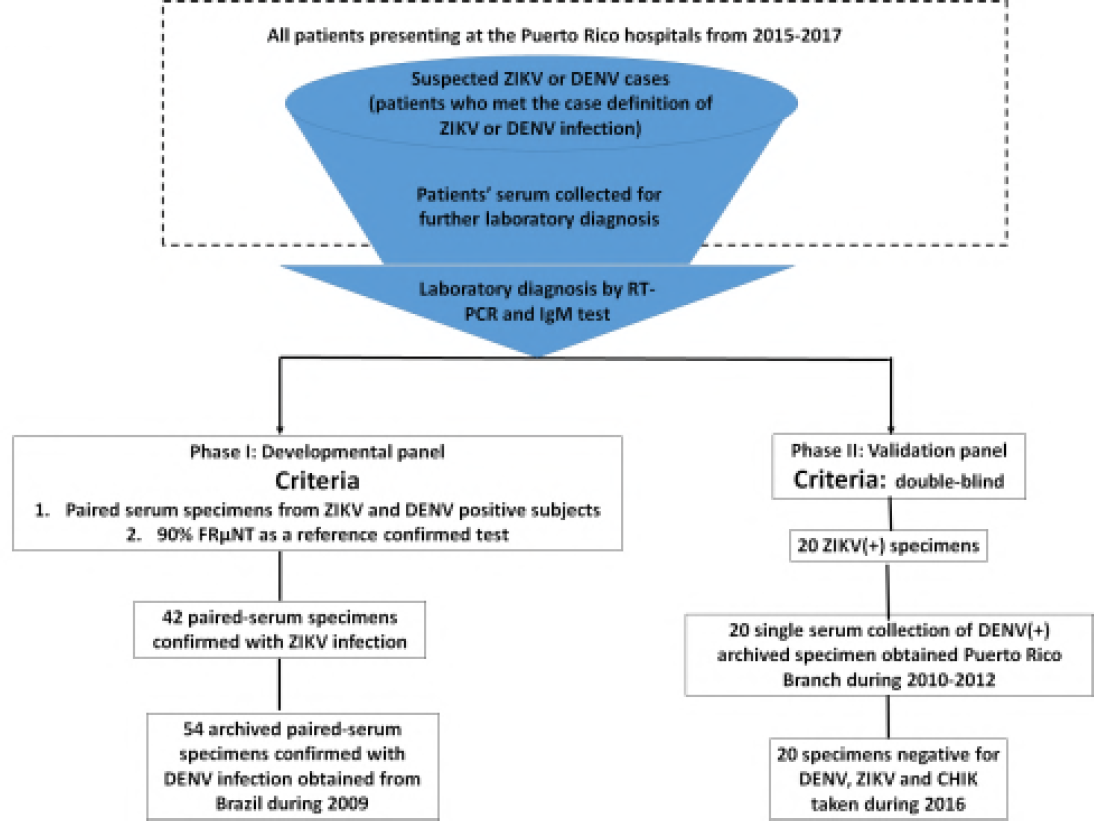
Flow chart of subject recruitment for the serum panels and case classification during Phase I and II. The source of ZIKV-infected serum specimens was all participants presenting at the Puerto Rico hospital from 2015-2017. Suspected cases of flavivirus infection were those with clinical symptoms matching the case reporting criteria defined by US-CDC and were admitted to the hospital for further diagnosis. All patients were classified as ZIKV or DENV infection if the results of ZIKV- or DENV-specific RT-PCR confirmed the diagnosis, respectively. Patients who tested negative by either ZIKV- or DENV-specific RT-PCR will be further subjected to an IgM test for the sera collected during a convalescent phase. Only the specimens that were ZIKV-specific RT-PCR positive in the acute phase and had seroconversion of IgM in the convalescent phase were included here. Those without a recorded RT-PCR or IgM laboratory results were excluded. The criteria of all ZIKV and DENV-confirmed specimens used in Phase I included paired sera collected during the acute and convalescent phase of illness and confirmation of disease status by FRμNT_90_ on ZIKV and DENV-2 in order to proper establishment of the cut-off value of the assay. The only criteria for the serum panel used in Phase II was double-blinded. All DENV positive specimens were collected during 2010-2012 during the time the paired specimen from the same patient was rare. The negative specimens were taken during 2016 and they are all negative for DENV, ZIKV, and Chikungunya virus (CHIK) in acute phase and negative for IgM in convalescent samples.

Detail characteristics of four groups of well-characterized, archived patient serum specimens used in Phase I were outlined in Table 1. To meet the outbreak reality when the paired specimens were difficult to obtain, the only criteria for the serum panel used in Phase II was double-blinded. All DENV positive specimens were collected during 2010-2012 when the paired specimen from the same patient was rare. The negative specimens were taken during 2016 and they are all negative for DENV, ZIKV, and Chikungunya virus (CHIK) in acute phase and negative for IgM in convalescent phase samples.

### Establishment of ZIKV NS1-MAC-ELISA

IgM antibody-capture ELISA has traditionally been used to selectively detect the IgM antibodies and to avoid the competition between IgM and IgG for a specific target antigen (such as prM/E containing flavi-VLP antigens). This is in contrast of using E or NS1 antigens for direct detection of anti-E or anti-NS1 antibodies in other studies.^18, 25, 26, 27^ The use of VLPs in MAC-ELISA has good sensitivity, safety, and acceptable specificity for determining a current flaviviral infection and was chosen here. Based on our previous publications^22^, depletion of anti-prM/E antibodies in advance is necessary to detect flavivirus-specific anti-NS1 antibodies using MAC-ELISA. To confirm if this is true for ZIKV infection, ZIKV-infected patient sera were added to pre-coated ELISA plates with or without depletion of prM/E antibodies. As shown in Fig 2A, depletion of anti-prM/E antibodies significantly enhanced the sensitivity of detecting anti-NS1 antibodies.

**Fig 2.**
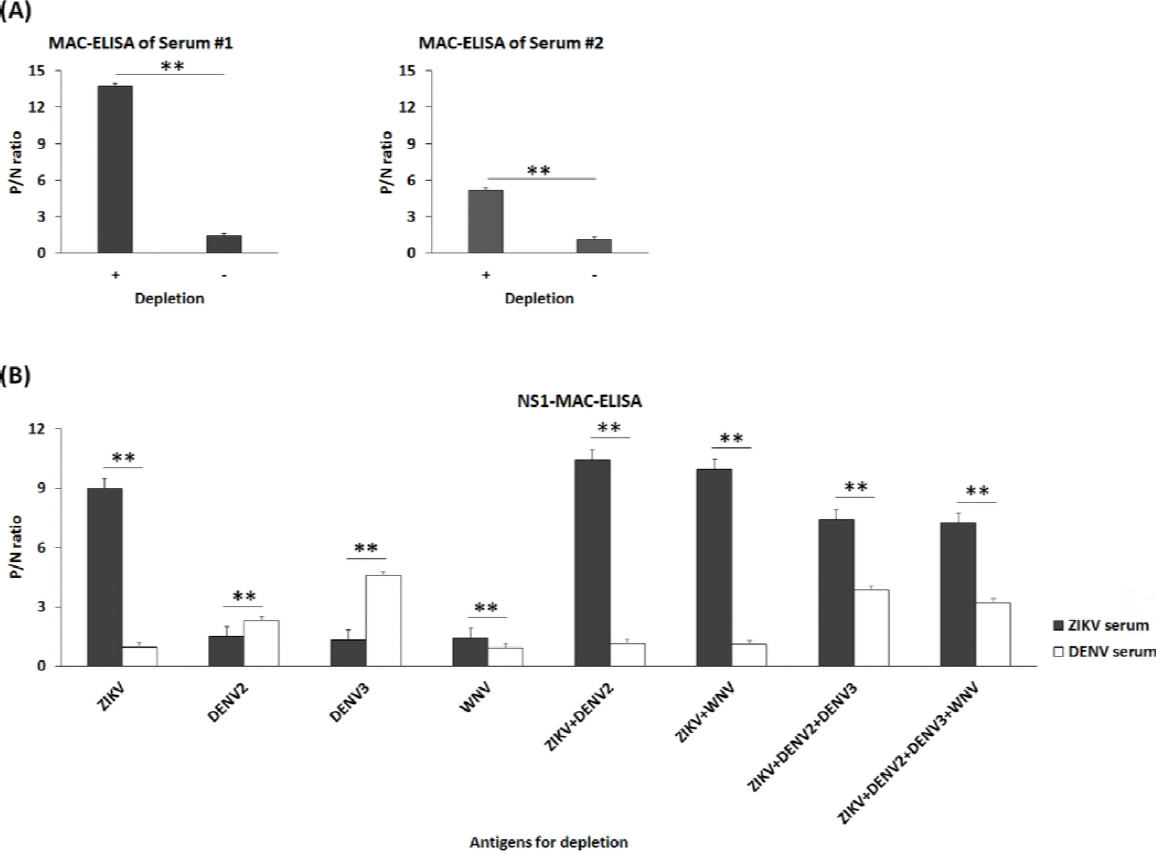
Analysis of the effect of depletion of anti-prM/E antibodies on NS1-MAC-ELISA using ZIKV VLP alone or in combination with VLPs from DENV-2, DENV-3 and WNV. (A) Values of P/N ratio of IgM and IgG from ZIKV-infected patient serum #1 (primary Zika infection 48B) and #2 (secondary Zika infection 45B) using ZIKV VLP alone for depletion and ZIKV-NS1 antigens for NS1-MAC-ELISA (+ on the *x* axis indicated with depletion and – indicated without depletion). (B) Values of P/N ratio of IgM and IgG from one ZIKV- (Black bar) or DENV-infected (white bar) patient serum using single, double, triple or quadruple VLP antigens of ZIKV, DENV-2, DENV-3, WNV for depletion. Normal human serum was used as a negative control to calculate the P/N ratio value by dividing the OD450 of ZIKV or DENV-confirmed patient serum with that of negative-control serum. All data were obtained from three independent experiments, and standard deviations are indicated. **p<0.0001.

Since ZIKV and DENV co-circulate in the same geographic location, we determined if a combination of multiple flavivirus VLPs could be required for depletion when the status of infection from the patient serum is unknown. As shown in Fig 2B, using a single serotype of DENV VLP could potentially result in a false negative result if infection from a different serotype of DENV occurred, which is consistent with our previous publication.^22^ Although using the combination of VLPs from ZIKV, DENV-2 and DENV-3 slightly decreased the P/N ratio from ZIKV serum panel, significantly increase the P/N ratio from DENV serum panel was noticed. Our results showed that the combination of VLPs from ZIKV, DENV-2, and DENV-3 was the best antigen combination to deplete the dominant flavivirus VLP-antibodies for the detection of NS1 antibodies against ZIKV- or DENV-infected patient serum specimens.

### Cross-reactivity of anti-prM/E and anti-NS1 antibodies

In order to compare the cross-reactivity of anti-prM/E and anti-NS1 antibodies, the proper cut-off values for VLP- and NS1-MAC-ELISAs were determined using a well-characterized control serum panel. Fig 3A shows the results of a ZIKV-VLP-MAC-ELISA for four different serum panels including antibodies to ZIKV, WNV, DENV and others (including other flaviviruses). Significant elevation of the P/N ratio in the convalescent phase sera was observed for patient antisera from both ZIKV and DENV infections. WNV and other anti-serum panels have little cross-reactivity with ZIKV-VLP, and were used to determine the cut-off values for the ZIKV-VLP-MAC-ELISA. The ROC analysis in Figure 3B, showing the curve of sensitivity vs 100-specificity, provides information on how strongly a given test result can be used to predict the likelihood of evidence of infection or non-infection based on P/N values from forty-two ZIKV patient sera and the WNV/other control serum panels (Fig 3B). The optimal cutoff values of P/N ratios for both ZIKV-VLP-MAC-ELISAs of acute and convalescent phase sera were set at 2.837 and 2.76, respectively (Fig. 3C). Similarly, the results of ZIKV-NS1-MAC-ELISAs for four different serum panels are shown in Fig 4A. The optimal cutoff values of P/N ratios for ZIKV-NS1-MAC-ELISAs of both acute and convalescent sera were set at 1.014 and 1.136, respectively (Fig. 4B and 4C).

**Fig 3.**
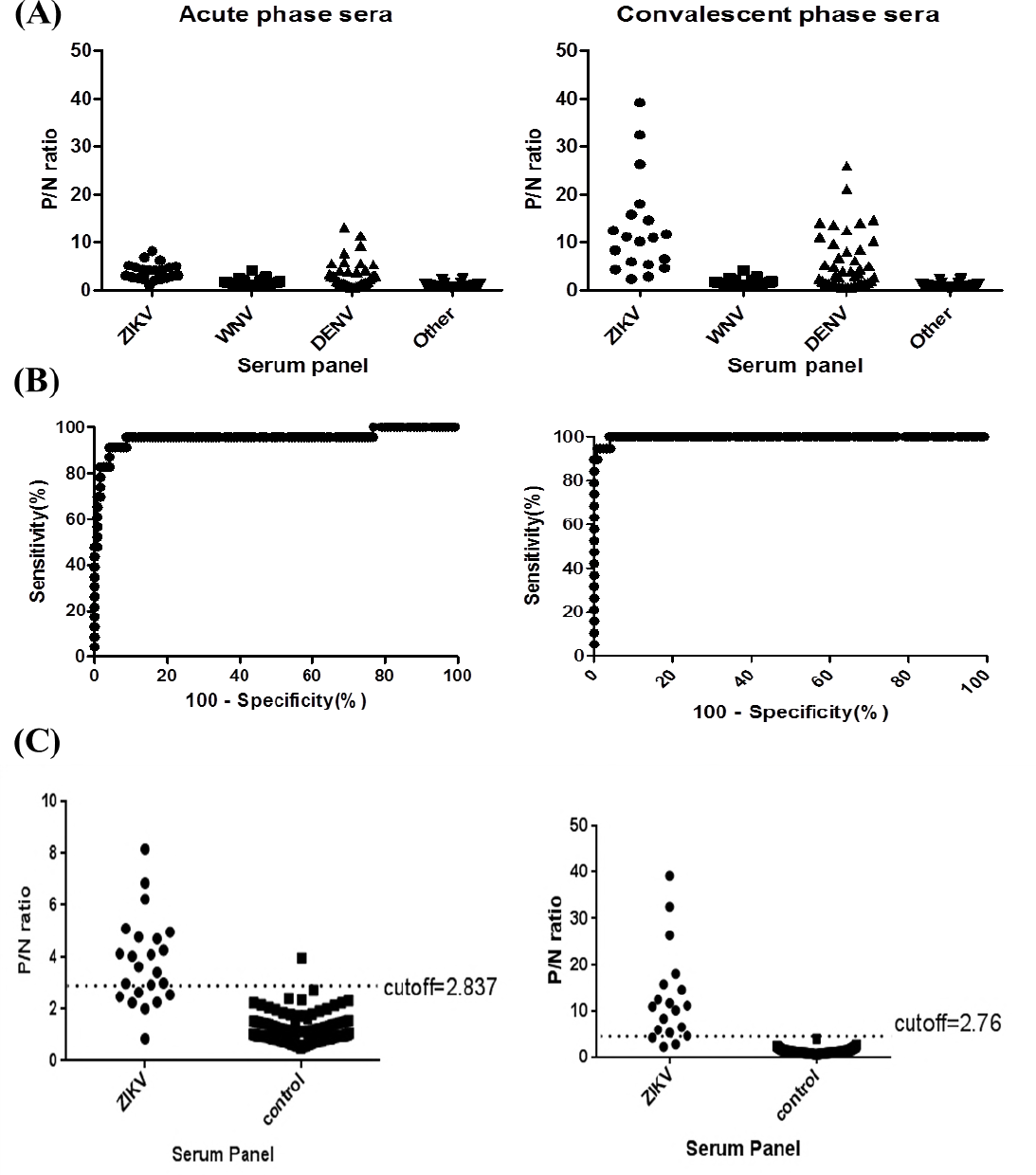
Distribution of P/N ratio of four groups of human patient sera and the determination of optimal cutoff P/N value of ZIKV-VLP-MAC-ELISA from acute (left panel) and convalescent (right panel) sera. (A) Values of P/N ratio for ZIKV, WNV DENV and other serum specimens. (B) The plot of sensitivity versus 100-specificity based on P/N values from 42 ZIKV-confirmed sera and 173 control serum panel. (C) Optimal cutoff value was determined by the magnitude of likelihood ratio positive (LR+) calculated by dividing sensitivity by 100-specificity.

**Fig 4.**
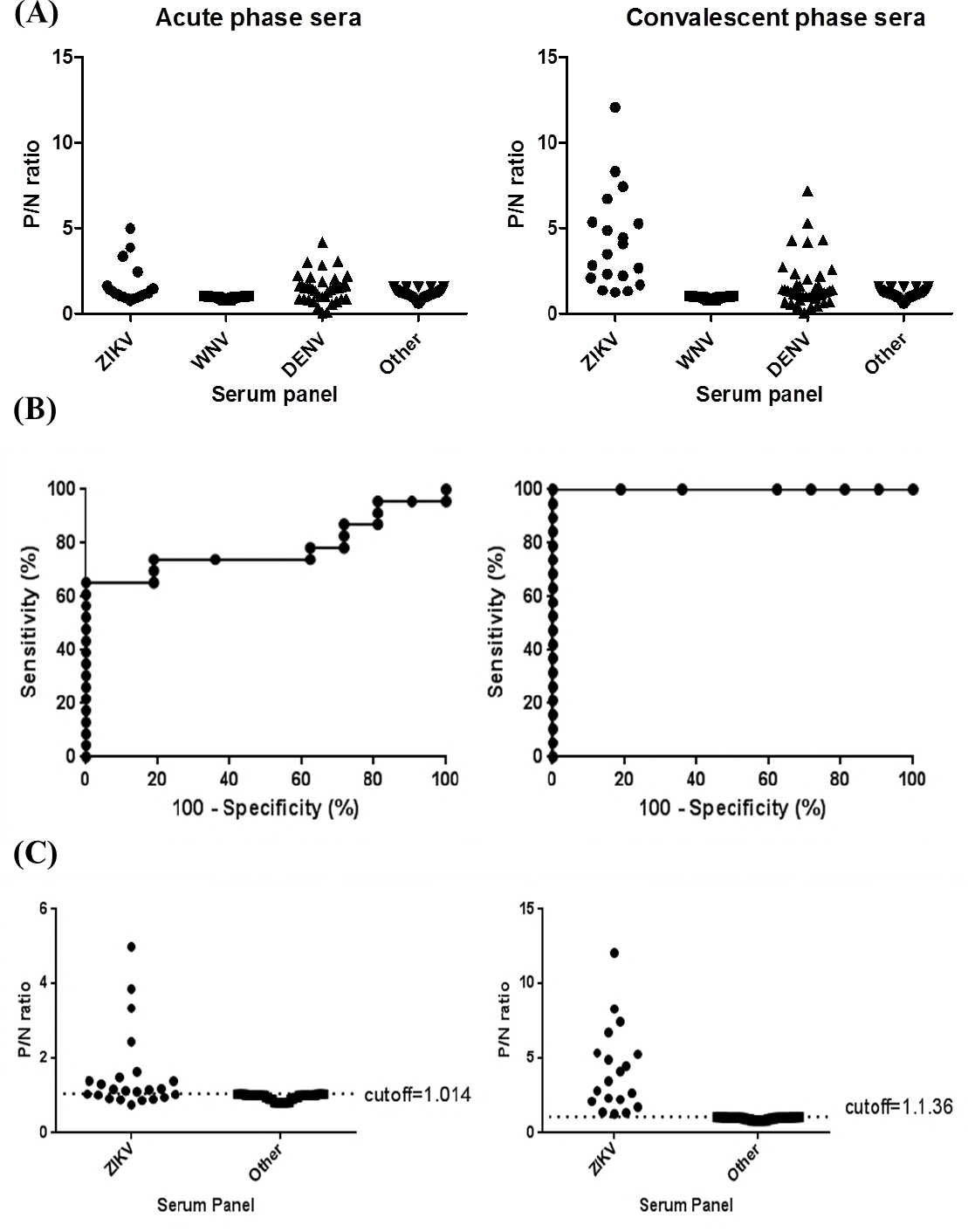
Distribution of P/N ratio of four groups of human patient sera and the determination of optimal cutoff P/N value of ZIKV-NS1-MAC-ELISA from acute (left panel) and convalescent (right panel) sera. (A) Values of P/N ratio for ZIKV, WNV DENV and other serum specimens. (B) The plot of sensitivity versus 100-specificity based on P/N values from 42 ZIKV-confirmed sera and 173 control serum panel. (C) Optimal cutoff value was determined by the magnitude of likelihood ratio positive (LR+) calculated by dividing sensitivity by 100-specificity.

Based on the cutoff, similar percentages of ZIKV acute (69.6% and 69.6%), and convalescent sera (94.7% and 100%) were positive for both ZIKV-VLP and NS1-MAC-ELISA, respectively (Table 2). However, significant numbers of the DENV panel were also positive to ZIKV VLP (63.6%) and NS1-MAC-ELISAs (72.7%). When ZIKV serum specimens were tested against DENV2 VLP and NS1 antigens, 95.7% were positive to VLP but only 8.7% were positive to NS1 from acute phase sera. On the contrary, 100% and 52.6% of the convalescent phase sera were positive for VLP and NS1 antigens of DENV-2, respectively. In summary, our VLP- and NS1-MAC-ELISAs have similar sensitivity detecting anti-prM/E and NS1 antibodies from ZIKV-infected patient sera. Although a significantly lower percentage of ZIKV patient sera was positive to NS1 antigens than VLP of DENV-2, no difference of cross-reactivity to ZIKV antigens was observed for DENV patient sera.

**Table 2.** Cross-reactivity of antibodies from four groups of human patient serum specimens against ZIKV and DENV-2 VLP and NS1 antigens.

| Serum panel |  | ZIKV-VLP-<br>MAC-ELISA | ZIKV-NS1-<br>MAC-ELISA | DENV2-VLP-<br>MAC-ELISA<sup>+</sup> | DENV2-NS1-<br>MAC-ELISA* |
| --- | --- | --- | --- | --- | --- |
| ZIKV (21) |  | 21/21(100%) | 21/21(100%) | 20/21(95.2%)* | 12/21(57.1%)* |
|  | Acute (23) | 16/23(69.6%) | 16/23(69.6%) | 22/23(95.7%)** | 2/23 (8.7%)** |
|  | Convalescent(19) | 18/19(94.7%) | 19/19(100%) | 19/19(100%*** | 10/19(52.6%*** |
| WNV (56) |  | 2/56 (3.6%) | 0/56 (0%) | NA | NA |
| DENV (44) |  | 28/44(63.6%) | 32/44(72.7%) | 42/44(95.5%) | 40/44(90.9%) |
|  | Acute (40) | 17/40(42.5%) | 25/40(62.5%) | 26/40(65.0%) | 23/40(57.5%) |
|  | Convalescent(42) | 23/42(54.8%) | 24/42(57.1%) | 40/42(95.2%) | 38/42(90.5%) |
| Other(76) |  | 0/76 (0%) | 0/76 (0%) | 0/76 (0%) | 2/76 (2.6%) |

NA: not available
<sup>+</sup>The cutoff of DENV-2 VLP or NS1-MAC-ELISA was based on our previous publication (??).
\*P=0.0042 with significance level at 0.05
\*\*p<0.0001 with significance level at 0.05
\*\*\*p=0.0007 with significance level at 0.05

### Using ZIKV/DENV2 ratio to differentiate between ZIKV and DENV infection

Previous studies suggested that a significant proportion of anti-E antibodies were GR antibodies that recognized the highly conserved fusion peptide during flavi virus infection.^24, 28^ To avoid the binding of such GR antibodies on the ZIKV VLP, a fusion peptide mutant ZIKV VLP (ZIKV FP-VLP) was generated for MAC-ELISA and the proper cutoff value was determined (Fig S1). A significant decrease of the cross-reactivity to ZIKV FP-VLP among the DENV serum panel was noticed; in addition, a similar positive proportion among the ZIKV serum panel was detected (Table S1). Therefore, our data suggested that FP-VLP, compared to WT-VLP as a diagnostic reagent, would be a more specific antigen for detecting anti-prM/E antibodies although there were still nearly 40% of DENV sera cross-reactive with ZIKV FP-VLP.

Further comparing the P/N ratios of the MAC-ELISA between ZIKV and DENV-2 FP-VLP, consistently higher values were observed when using homologous antigens; that is, for a ZIKV infection, higher values of the P/N ratio was observed for ZIKV FP-VLP than for the use of DENV-2 FP-VLP (Fig 5). Similar results were also observed for the NS1-MAC-ELISA. Therefore, an algorithm of serological diagnosis to differentiate between ZIKV and DENV infection was developed in this study (Fig 6).

**Fig 5.**
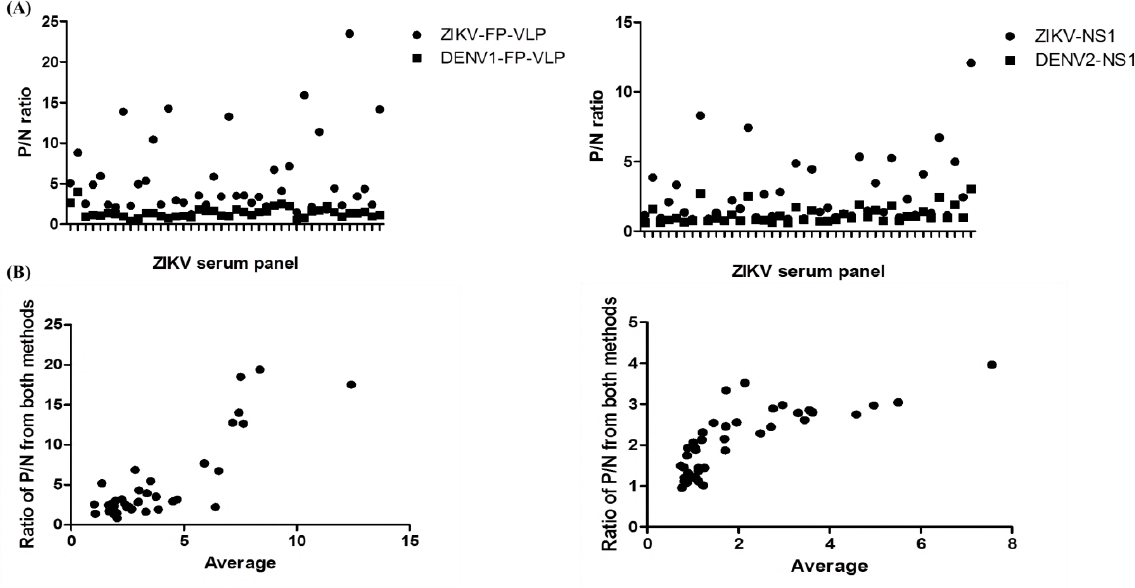
P/N values (A) and Blant-Altman plots (B) of FP-VLP- and NS1-MAC-ELISA from 42 ZIKV-patient’s serum specimens.

**Fig 6.**
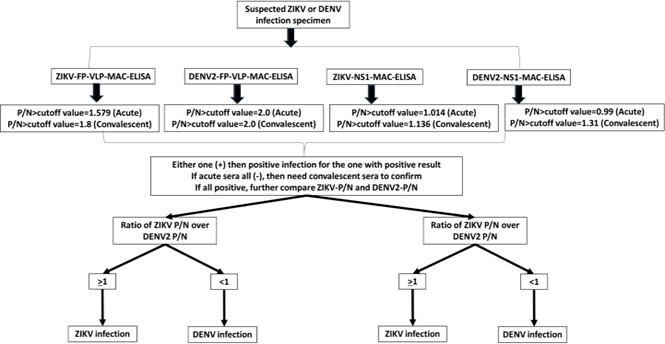
Algorithm of differentiating ZIKV and DENV infection by VLP-MAC-ELISA and NS1-MAC-ELISA.

### Validation of the algorithm

The prospectively collected validation serum panel was provided to the investigator and blind tested by VLP-MAC-ELISA and NS1-MAC-ELISA using FP-VLP and NS1 from ZIKV and DENV-2 and the results were interpreted based on the developed algorithm in Fig 6. Twenty (100%) of the ZIKV-confirmed sera were classified as ZIKV infection by FP-VLP-MAC-ELISA and 80% by NS1-MAC-ELISA. For DENV-confirmed specimens, 75% and 100% were classified as DENV infection by FP-VLP-MAC-ELISA and NS1-MAC-ELISA, respectively. Fifteen percent of the negative specimens were falsely classified as positive by FP-VLP-MAC-ELISA and 10% by NS1-MAC-ELISA. Overall, the sensitivity and specificity of FP-VLP-MAC-ELISA and NS1-MAC-ELISA based on the algorithm was higher than 80% with no statistical significance, although slightly lower sensitivity (75%) of FP-VLP-MAC-ELISA in classifying DENV infection (Table S2) was observed. The overall PPV of both assays for diagnosis of ZIKV or DENV infection demonstrates no statistical significance.

## Discussion

So far, no publications have evaluated anti-E or anti-NS1 antibodies against DENV or ZIKV simultaneously using well-archived human serum samples. In this study, we comprehensively evaluated the cross-reactivity of anti-prM/E antibodies induced by ZIKV infection using wild-type and FP-mutated VLP antigens from ZIKV and DENV-2. The test results were compared to the prM/E antibody-depleted NS1 MAC-ELISA. Using ZIKV-FP-VLP significantly reduced the observed cross-reactivity for a DENV patient serum panel, compared to using wild-type ZIKV VLP. Although ZIKV-NS1-MAC-ELISA is more specific in determining ZIKV infection, we still observed 57.1% cross-reactivity for ZIKV infection and 90.9% for DENV infection, which is consistent with a previous publication.^25^ Using a combination of a ZIKV/DENV-2 ratio from VLP- and NS1-MAC-ELISAs, we successfully differentiated between ZIKV and DENV infection with 90-100% accuracy. Thus, we have demonstrated a testing algorithm for differentiating ZIKV and DENV infections that can be applied in dengue- and/or other flavivirus-endemic regions where most patients have had a previous flavivirus infection.

Using rigorous evaluation, our study compared the cross-reactivity of anti-prM/E and anti-NS1 antibodies across different sero-complexes, including ZIKV, DENV, WNV and others, by using well-characterized, archived serum specimens. The overall cross-reactivity of anti-NS1 antibodies induced by ZIKV infection was significantly lower than anti-prM/E antibodies, possibly due to the difference in electrostatic surface potential of NS1.^8, 29, 30^ However, we did not observe any significant differences in cross-reactivity between VLP- and NS1-MAC-ELISAs for DENV infection. The results of highly cross-reactive anti-prM/E antibodies were consistent with the focus-reduction micro-neutralization test (FRμNT) results (Table S3). The majority of ZIKV-infected patients had prior DENV infection as suggested by FRμNT titers greater than 10 against DENV-2 in acute serum specimens. During secondary ZIKV infection, the FRμNT titer in the convalescent sera showed at least 4-fold increase against both ZIKV and DENV. The majority of cross-reactive anti-prM/E antibodies during flaviviral infection are GR (4G-2 like) antibodies recognizing the FP, with the potential to enhance viral infection and induce low-to-moderate neutralizing activity.^7^ The current CDC guideline suggests that a supplemental neutralization test be performed for all specimens positive by a ZIKV MAC-ELISA due to the possibility of antibody cross-reactivity.^20^ Our current results further suggest that a 90% FRuNT should be performed to distinguish between ZIKV and DENV infection.

Flaviviruses have been traditionally subdivided into different serocomplexes, comprised of members that are cross-neutralized by polyclonal sera.^10^ Such sero-classification was correlated with the similarity of amino acid sequence of prM/E.^7^ ZIKV, clustered with Spondweni virus and shows an intermediate position with viruses from JEV and DENV serocomplexes in the phylogenetic tree (based on complete genome, E, or NS1 gene sequences). The overall picture of flavivirus serocomplexes indicate that cross-neutralization with polyclonal sera is usually lost when the amino acid sequence divergence of E is more than 50%.^7^ Therefore, ZIKV together with the viruses from the Spondweni virus group could form an independent serocomplex, which serves as the basis of using the ratio of ZIKV/DENV IgM antibodies for a differential diagnosis. A similar concept could also be applied to anti-NS1 antibodies, which share a similar amino acid sequence divergence with viruses from different serocomplexes. The use of FP-VLP in this study has several advantages, including the avoidance of a pre-depletion step prior to detecting anti-NS1 antibodies, and the reduced binding of cross-reactive fusion-loop antibodies, which significantly enhance the specificity and accuracy of using a ZIKV/DENV ratio in differentiating ZIKV and DENV infections.

Our study revealed a similar percentage of ZIKV acute (69.6% and 69.6%), and convalescent sera (94.7% and 100%) were positive for both ZIKV-VLP and NS1-MAC-ELISA, respectively. Previous publications suggested a lower sensitivity of detecting anti-NS1 antibodies, possibly due to their relatively low abundance compared to anti-prM/E antibody in human sera.^15, 22^ Our study showed that depletion of anti-prM/E antibodies enhances the sensitivity of NS1 antibody detection. Based on the testing algorithm developed in this study, FP-VLP-MAC-ELISA and NS1-MAC-ELISA had similar PPV and NPV (Table 2). By combining both assays, the accuracy of sero-diagnosis can reach up to 95% (57/60) using the classification guideline provided in Table S4. Considering the severe outcome of congenital Zika syndrome, three false positive specimens (5%), misclassified as ZIKV infections, may be acceptable (Table S5). The important limitations of the current study is the small sample size of the validation serum panel and the generalizability to a more complex serum panels such as subjects with prior exposure to St. Louis Encephalitis virus, Japanese encephalitis virus, Powassan virus, and yellow fever virus. In summary, the current study successfully develops a novel approach to accurately differentiate ZIKV and DENV infections for evidence-based public health intervention.

## Contributors

MTW and BSD performed all the construction of recombinant NS1 and VLP expression plasmids as well as rabbit serum production. DYC and GJC designed the experiments and wrote the manuscript. DYC performed all the ELISA and statistical analysis. FAM and JLM characterized all ZIKV and DENV serum panels. All authors reviewed the draft, had critical input, and reviewed the final submission.

## Acknowledgements

DYC was supported by MOST oversea short-term fellowship in CDC to conduct this study. We are thankful Ann Hunt and Ann Powers for scientific comments and English editing.

